# New races with wider virulence indicate local evolution of *Puccinia striiformis* f. sp. *tritici* in South America

**DOI:** 10.1101/2023.10.23.563651

**Authors:** V. Riella, J. Rodriguez-Algaba, R. García, F. Pereira, P. Silva, M.S. Hovmøller, S. Germán

## Abstract

Wheat yellow (stripe) rust, caused by *Puccinia striiformis* f. sp. *tritici* (*Pst*), is one of the most devastating diseases of wheat worldwide. *Pst* populations are composed of multiple genetic groups, each carrying one or more races characterized by different avirulence/virulence combinations. Since the severe epidemics in 2017, yellow rust has become the most economically important wheat foliar disease in Uruguay. Evolution of virulence was investigated based on genotyping and race typing of a representative set of 27 *Pst* isolates collected from wheat fields in Uruguay between 2017 and 2021. Three genetic groups were identified, i.e., *PstS7*, *PstS10* and *PstS13*, the latter being the most prevalent. Two races previously reported in Europe, Warrior (*PstS7*) and Benchmark (*Pst*S10), were detected in four and two isolates, respectively. A third race known as Triticale2015 (*PstS13*), first detected in Europe in 2015 and in Argentina in 2017, was detected at several locations. Additional virulence to *Yr3, Yr17*, *Yr25*, *Yr27* or *Yr32* was detected in three new race variants within *PstS13*. The identification of these new races, which have not been reported outside South America, provides strong evidence of the local evolution of virulence in *Pst* during the recent epidemic years.

## Introduction

Wheat yellow (stripe) rust, caused by *Puccinia striiformis* f. sp. *tritici* (*Pst*), is one of the most devastating diseases of wheat worldwide (Chen et al. 2014; Wellings 2011; Stubbs 1985; Beddow et al. 2015). *Pst* is favored by relatively low temperatures of 10-15°C and causes very significant grain yield losses in highly susceptible cultivars (Chen 2005; Carmona et al. 2019; Roelfs et al. 1992). Historically, *Pst* has mainly been a problem in cool climates, however, in recent years, the pathogen has gained tolerance to higher temperatures and become an increasing problem in areas normally considered too warm for *Pst* establishment (Wellings 2007; Milus et al. 2009). Moreover, *Pst* epidemics originating from distant geographical areas have been reported, either as an incursion to new regions where it was previously absent or as a re-emergence of new races with increased aggressiveness (Bahri et al. 2009; Hovmøller et al. 2023b). As a consequence, *Pst* epidemics have been an increasing problem threatening global wheat production (Boshoff et al. 2002; Ali et al. 2014; Hovmøller et al. 2016).

*Pst* was detected for the first time in Uruguay and Argentina in 1929 (Rudolf and Job 1931). During 1929 and 1930, *Pst* caused widespread epidemics through most of the Southern Cone region, causing extremely high yield losses (Boerger 1934; Vallega 1938). From its first detection until 2016, *Pst* occurred sporadically, rarely reaching epidemic levels in Uruguay (Germán and Caffarel 1999; Germán et al. 2007, 2018). Since 2017, *Pst* has caused generalized epidemics in Uruguay and Argentina (Germán et al. 2018, 2021; Carmona and Sautua 2018; Campos 2020). *Pst* is currently the most economically important wheat foliar disease, requiring the highest number of fungicide applications. Locally, on susceptible and moderately susceptible cultivars, farmers typically apply two fungicide applications each growing season to obtain adequate disease control. The most susceptible cultivar to *Pst* included in the National Cultivar Evaluation trials in Uruguay, had grain yield losses ranging between 71% and 82% in 2017 (Germán et al. 2018). In Argentina, races overcame many of the major resistance genes in germplasm locally adapted (Carmona et al. 2019). Since 2020, more than 50% of the wheat cultivars sown in Uruguay appeared as either susceptible or moderately susceptible to *Pst*.

Mutations and subsequent selection are considered the main driving forces that generates new races with virulence to the deployed host resistance genes (de Vallavieille-Pope et al. 2012; Hovmøller and Justesen 2007). *Pst* isolates undergo step-wise mutations, generating isolates that differ in virulence (Park 2015). Sexual recombination is another mechanism that may generate variability in *Pst*, which has been reported under experimental conditions involving the alternate host *Berberis spp.* (Rodriguez-Algaba et al. 2014, 2020), however, rarely reported under natural conditions. As these secondary hosts are not reported in Uruguay, sexual recombination is thought to have no impact as a source of new variability locally, leading to the assumption of a strongly clonal *Pst* population in Uruguay as reported for other regions (Ding et al. 2021; Hovmøller et al. 2002).

*Pst* races are characterized phenotypically through their avirulence/virulence pattern to *Yr* resistance genes. Additionally, the *Pst* genetic diversity can be studied using microsatellite molecular markers to assign the different *Pst* isolates to different genetic groups (Bai et al. 2021; Sharma-Poudyal et al. 2020; Walter et al. 2016; Ali et al. 2017), allowing a better understanding of the evolutionary and dispersal dynamics of the pathogen (Thach et al. 2016; Ali et al. 2014; Ding et al. 2021).

Recent *Pst* epidemics worldwide and the spread of epidemics to new areas, where the disease was previously nonsignificant, make it urgent to understand the evolution of *Pst* races, their spread and establishment. Knowledge of the evolution of the pathogen virulence is a key factor in determining the best strategy to breed locally adapted cultivars with effective and durable resistance to *Pst*. In the present study, we examined the population structure of *Pst* in Uruguay based on samples collected from wheat fields between 2017 and 2021. The *Pst* isolates were assigned to already defined genetic groups and race typed to study local evolution of virulence within these groups, which may have strong implications for prevention and control of rust diseases in wheat.

## Material and methods

A set of representative 27 isolates were selected from the Uruguayan collection of *Pst* samples maintained at INIA-La Estanzuela (Colonia, Uruguay). The collection was composed of 128 isolates, recovered from a total of 260 samples collected from spring bread wheat (*Triticum aestivum*) at different locations in the principal wheat production area in Uruguay between 2017 and 2021. Isolates were selected based on year, location, and preliminary race typing information generated locally. From the 27 selected samples, 4 samples were collected in 2017, 5 in 2018, 2 in 2019, 11 in 2020 and 5 in 2021 (Supplementary material 1).

### Sample recovery and race typing

The original samples, preserved at 5°C, were used to inoculate seedlings of susceptible wheat cultivar Morocco. Infected leaves were collected separately and sent to the Global Rust Reference Center (GRRC) in Denmark following their standard practice recommendations for sample shipment. For isolate testing at the GRRC, the protocol described in Hovmøller et al. (2017) was used. Leaf segments were placed in Petri dishes with moist filter paper and incubated between 24h to 48h to promote spore production.

Seedlings of wheat cultivars Morocco and Morocco/*Lr*19 were used for multiplication of spores. Seedlings were grown in pots, 12–15 plants per pot, and treated with 5ml of 0.5% Antergon MH180 (Nordisk Alkali, Randers, Denmark) to regulate plant growth and enhance spore production. Leaves with newly emerged urediniospores were gently rubbed on the wheat seedlings. Inoculated seedlings were misted with water and incubated in a dew chamber at 10–12°C in darkness for 24h, then transferred to spore-proof greenhouse cabins at 17°C during the day/12°C during the night with a 16h photoperiod of natural light and supplemental sodium light (100μmol/s/m) and 8h darkness, relative humidity of 70–80%. Pots were covered with cellophane bags (Helmut Schmidt Ver-packungsfolien GmbH, Königswinter, Germany) prior to sporulation to prevent cross-contamination among isolates. Spores were harvested by shaking the plants inside the cellophane bag, and then transferred to cryovials. These were then dried in a desiccator for approximately three days and preserved at -80°C until further use.

All isolates were race typed using the current standard set of 24 wheat differential lines (Table 1). A supplementary test was carried out including additional lines for further confirmation of specific race variants. Previously characterized reference isolates of the genetic groups detected for the Uruguayan isolates were included as controls.

**Table 1.** Wheat differential lines used for race typing of *Puccinia striiformis* f. sp. *tritici* isolates.

| Differential line | Yellow rust resistance genes (*Yr*) | GRRC Standard set | Supplementary test |
| --- | --- | --- | --- |
| Chinese 166 | *1* | x |  |
| Kalyansona | 2, + | x | x |
| Vilmorin 23 | 3, + | x | x |
| Hybrid 46 | 4, + | x |  |
| Suwon | *Su* | x |  |
| Heines Kolben | 6, + | x |  |
| Avocet *Yr6* | 6, *AvS* | x |  |
| Lee | 7, + | x |  |
| Avocet *Yr8* | 8 | x |  |
| Avocet *Yr9* | 9, *AvS* | x |  |
| Moro | 10 | x |  |
| Cortez | 15 | x |  |
| VPM1 | 17, + | x | x |
| Avocet *Yr17* | 17, *AvS* | x | x |
| TP 981 | 25, + | x | x |
| Spaldings Prolific | *Sp*, 25, + | x | x |
| Opata | 27, 18, + | x | x |
| Avocet *Yr32* | 32, *AvS* | x | x |
| Avocet S | *AvS* | x | x |
| Strubes Dickkopf | *Sd*, 25, + |  | x |
| Heines VII | 2, 25, + |  | x |
| Carstens V | 32, 25, + |  | x |
| Avocet *Yr27* | 27, *AvS* |  | x |
| Heines Peko | 2, 6, 25, + |  | x |
| Nord Desprez | 3 |  | x |
| Ambition | *Amb* | x |  |
| Benchmark | unknown | x |  |
| Kalmar | unknown | x |  |
| Nemo | unknown | x |  |

Ten to twelve seeds of each wheat genotype were sown in 7×7×8cm pots with a 1:1 Pindstrup Substrate peat mix containing slow-release plant nutrients (Pindstrup Mosebrug A/S, Ryomgaard Denmark). Differential sets with one pot of each wheat genotype were placed in a tray and gown in spore-proof cabins in the greenhouse at 17°C day/12°C night temperature regime and inoculated approximately 12 days after sowing when the second leaf was half unfolded. Urediniospores were retrieved from -80°C and used for inoculation after heat shock treatment in a water bath at 40–42°C for 2min. Approximately 25mg of spores were suspended in 3ml engineered fluid 3M™ Novec™ 7100, (3M, St. Paul, MN, USA) and gently mixed. Wheat seedlings were spray inoculated using an airbrush spray gun (standard class, Revell GmbH, Bünde, Germany) in a laboratory fume hood. Seedlings were subsequently sprayed with mist water, incubated in a dew chamber, and transferred to the greenhouse under the same conditions described above.

Infection type (IT) was scored on individual plants/leaves after 15–17 days. The first and the second leaves were scored separately using a 0–9 scale (McNeal et al. 1971), where scores between 7 and 9 indicated compatibility (virulence) and scores equal to or below 6 indicated incompatibility (avirulence).

### Genotypic characterization of isolates

The genotyping of isolates was based on 19 Simple Sequence Repeat (SSR) markers according to Rodriguez-Algaba et al. (2017). This allowed a genetic grouping based on genotypic similarities, i.e., minor (or no) allele differentiation among individuals within a group, and major differences between groups following the principles of Ali et al. (2017). Allele sizes detected for individual isolates and associated genetic groups are presented in Supplementary material 2.

## Results

### *Pst* genetic groups

The 27 isolates were assigned to one of the previously reported genetic groups if their genetic profiles for the set of 19 SSRs matched the reference profile (Hovmøller et al. 2016; Ali et al. 2017). Three different *Pst* genetic groups were detected in Uruguay (Supplementary material 2). *PstS13* was the most prevalent group with 78% of the isolates analyzed, followed by *PstS7* with 15% and *PstS10* with 8% (Table 2). The *PstS13* group was identified in samples collected from all sampling years, *PstS7* was found for the first time in 2018 and then in 2020 and 2021, and *PstS10* was only detected in 2020.

**Table 2.** Number of isolates of different genetic groups and virulence phenotypes for samples collected in Uruguay during 2017-2021.

| Genetic group | Race | Virulence phenotype | 2017 | 2018 | 2019 | 2020 | 2021 |
| --- | --- | --- | --- | --- | --- | --- | --- |
| *PstS7* | Warrior | 1,2,3,4,6,7,-,9,-,-,17,25,-,32,*Sp*,*AvS*,*Amb* |  | 1 |  | 2 | 1 |
| *PstS10* | Benchmark | 1,2,3,4,6,7,-,9,-,-,17,25,-,32,*Sp*,*AvS*,- |  |  |  | 2 |  |
| *PstS13* | Triticale2015 **a** | -,2,-,6,7,8,9,-,-,-,-,-,AvS,- | 4 | 4 |  |  |  |
|  | **b** | -,2,-,6,7,8,9,-,-,17,-,-,32,-,AvS,- |  |  | 2 | 6 | 3 |
|  | **c** | -,2,-,6,7,8,9,-,-,17,-,27,32,-,AvS,- |  |  |  | 1 |  |
|  | **d** | -,2,3,-,6,7,8,9,-,-,17,25,-,32,-,AvS,- |  |  |  |  | 1 |

Note. Virulence phenotype numbers indicate virulence to yellow rust resistance genes:
*Yr1*, *Yr2*, *Yr3*, *Yr4*, *Yr6*, *Yr7*, *Yr8*, *Yr9*, *Yr10*, *Yr15*, *Yr17*, *Yr25*, *Yr27*, *Yr32*, Spalding
Prolific (*Sp*), Avocet S (*AvS*) and Ambition (*Amb*), respectively. “-” indicates avirulence.
Letters **a-d** indicate different virulence phenotypes within genetic group *PstS13*.
Note. All isolates belonging to genetic group *PstS10* were virulent on differential line
Benchmark.

### Race typing

The virulence phenotype of the 27 isolates and their corresponding genetic group are presented in Table 2. The primary results of the phenotyped isolates are presented in Supplementary material 3. Four isolates with virulence on *Yr1, Yr2, Yr3, Yr4, Yr6, Yr7, Yr9, Yr17, Yr25, Yr32, YrSp, YrAvS,* and *YrAmb,* shared virulence phenotype with a reference isolate of the “Warrior” race (*PstS7*). Two isolates were assigned to the “Benchmark” race (*PstS10*), which is characterized by avirulence to cultivar Ambition and virulence on cultivar Benchmark. Four additional closely related virulence phenotypes were detected among the isolates belonging to *PstS13* (Figure 1). The first race within this group, detected in 2017 and 2018, was virulent on *Yr2, Yr6, Yr7, Yr8, Yr9* and *YrAvS* (Figure 1-a), which corresponded to the original Triticale2015 race. The supplementary virulence test allowed the verification of three new race variants (Supplementary material 4). One variant, first detected in 2019, had gained virulence on *Yr17* and *Yr32*; a second variant, collected in 2020, had additional virulence on *Yr27* and a third variant, collected in 2021, was virulent on *Yr3*, *Yr17*, *Yr25* and *Yr32,* in addition to the virulence detected in the original Triticale2015 race.

**Figure 1.**
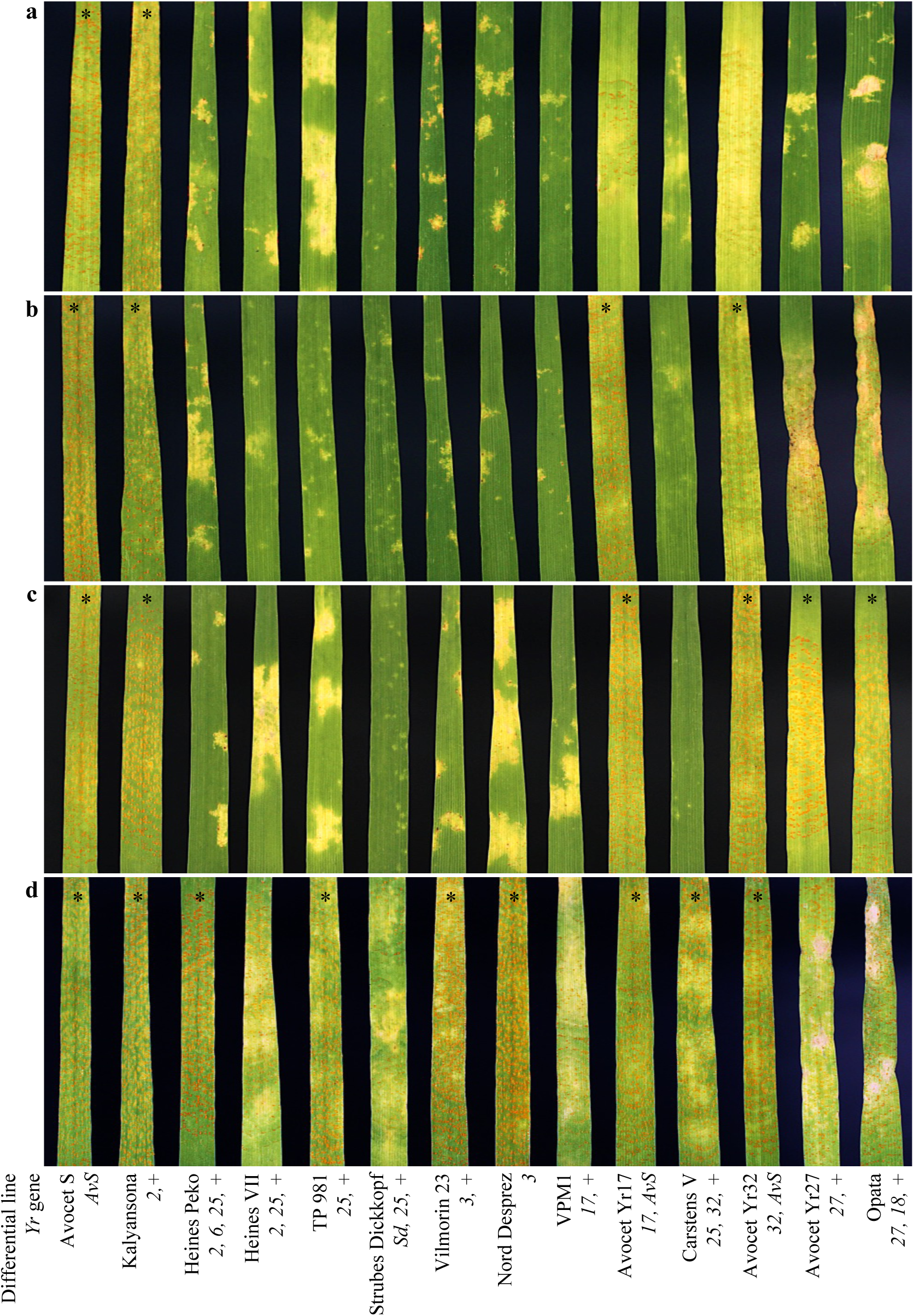
*Puccinia striiformis* f. sp. *tritici* infections on differential lines for the four different races (**a**-**d**) within Triticale2015 race (*PstS13* genetic group). Original Triticale2015 race **(a)**, races with additional virulence to *Yr17* and *Yr27* (**b**); to *Yr17*, *Yr27* and *Yr32* (**c**), and *Yr3*, *Yr17*, *Yr25* and *Yr32* (**d**), respectively. *: Compatible interaction (infection type 7-9).

## Discussion

A set of 27 recovered isolates from the Uruguayan *Pst* collection, sampled in Uruguay between 2017 and 2021 was analyzed. The set was considered representative of local *Pst* population based on collection year, location and preliminary race typing. The virulence and genotypic profiles allowed assignment to races within *PstS7*, *PstS10* and *PstS13* genetic groups, respectively, some of which previously reported as responsible for severe epidemics in the last decade (Beddow et al. 2015; Sørensen et al. 2014; Hovmøller et al. 2016; Ali et al. 2017). Three races previously described in Europe within each genetic group, were detected in this study, i.e., Warrior (*PstS7*), Benchmark (*PstS10*) and Triticale2015 (*PstS13*). Triticale2015, the original *PstS13* race, first detected in Europe in 2015 (Hovmøller et al. 2018) and in Argentina in 2017 (Carmona et al. 2019), was detected in several isolates. Additionally, three new race variants within the *PstS13* genetic group were also identified. These results provide strong evidence not only of the establishment of the disease in regions where *Pst* was not previously widespread, but also revealed the emergence of new race variants with wider virulence indicating the existence of local evolution of *Pst*.

The appearance in the Southern Cone of America of *Pst* genetic groups previously reported in Europe may be associated with human activities (Zadoks 1961; Stubbs 1985; Brown and Hovmøller 2002; Wellings 2007; Ali et al. 2014) and/or long distance wind dispersal (Zadoks 1961; Brown and Hovmøller 2002). Argentina and Uruguay are located in the same rust epidemiological region (Rajaram and Campos 1974), where there are no geographical barriers for urediniospores dispersal, which likely explains the almost simultaneous development of severe epidemics in both countries (Campos et al. 2016; Carmona et al. 2019; Germán et al. 2018). More recently *Pst* was detected in Paraguay for the first time (Fernández-Gamarra et al. 2023) and in Brazil. Before 2017, *Pst* did not cause severe epidemics in the epidemiological zone East of the Andes as the pathogen survived during the summer distant from the wheat crop regions (Germán et al. 2007). Since 2017, a relevant epidemiological change associated with the presence of *Pst* in Uruguay is the likely increased oversummering capacity of the pathogen. In fact, oversummering of the pathogen has been observed in the Argentinian wheat crop area, which allows primary inoculum to infect crops earlier and thus cause severe epidemics not only in Argentina but also in Uruguay (Germán et al. 2018; Silva et al. 2023).

Among the three genetic groups detected in this study, *PstS7* and *PstS10* had the lowest prevalence. Since 2011 and 2012, both genetic groups have been associated with important epidemics with substantial economic losses in various parts of the world (Hovmøller et al. 2016; Hubbard et al. 2015). In Europe, new races from *PstS7* and *PstS10* genetic groups have been detected since 2011 and have replaced the previous pathogen population (Sørensen et al. 2014; Ali et al. 2017). So far, only one race has been described within the *PstS7* genetic group (Warrior race), which has spread from Europe to North Africa (Hovmøller et al. 2016) and South America (Hovmøller et al. 2023a). *PstS10* has been the most prevalent genetic group in Europe from 2013 to 2022. One race has been prevalent within this genetic group, but some variants virulent to widely grown cultivars have been described (Hovmøller et al. 2020). In 2018, it was also detected in Australia (Ding et al. 2021; Park et al. 2020). Up to now, it has not yet been possible to differentiate the new race variants within *PstS10* by molecular techniques nor by standard wheat differential lines (Hovmøller et al. 2022). In terms of virulence, the races within *PstS10* are similar to the Warrior race (*PstS7*), except for their avirulence/virulence to the European wheat cultivars Warrior and Ambition (Hovmøller et al. 2020).

In this study, *PstS13* was the most prevalent genetic group. *PstS13* was first reported in Europe in 2015, mainly affecting triticale and durum wheat (Hovmøller et al. 2018). A single race within this genetic group is still prevalent in Europe although local variants have been observed (Hovmøller et al. 2018). The same genetic group caused severe epidemics on durum and bread wheat in Italy in 2017 (Hovmøller et al. 2018). A new *Yr10* virulent variant was detected in Poland in 2019 and in Germany in 2020 (Hovmøller et al. 2021). In Australia, *PstS13* was detected in 2018 and has become one of the most widespread genetic groups (Ding et al. 2021; Park et al. 2020). In South America, *PstS13* was reported as the prevalent genetic group in Argentina in 2017 and 2018 (Carmona et al. 2019; Hovmøller et al. 2018, 2019). Samples collected in Chile during 2018 were also assigned to this genetic group (Hovmøller et al. 2019). All samples from Argentina and Chile showed the same virulence phenotype as the original *PstS13* race reported in Denmark (Hovmøller et al. 2019, 2020). More recently, *PstS13* spread to Paraguay (Fernández-Gamarra et al. 2023). Since *PstS13* has been the most prevalent genetic group in South America accounting for severe *Pst* epidemics in the region, thus new virulence variants could be expected based on previously reported high mutation rates in yellow/stripe rust (Hovmøller and Justesen 2007).

Here we report new race variants with additional virulence within the *PstS13* genetic group. The emergence of new *Pst* races within the same genetic group is probably due to mutation and subsequent selection of races with virulence to the deployed host resistance genes (de Vallavieille-Pope et al. 2012; Hovmøller and Justesen 2007). Through this mechanism, *Pst* isolates acquire the ability to avoid recognition by resistance genes in host plants resulting in step-wise mutations (Hovmøller et al. 2002). This was already observed in Australia, where the sexual cycle of *Pst* does not occur, and new races usually differ from existing races by virulence on a single resistance gene (Wellings and McIntosh 1990; Park 2015; Ding et al. 2021). The emergence of new races may be related to the presence of the corresponding resistance genes in wheat cultivars used locally. One possible pathway to explain the observed pathogen evolution in Uruguay starts with the original *PstS13* detected in samples from 2017 and 2018 (*PstS13*a, Table 2). *PstS13*b, detected in 2019, gained virulence on *Yr17* and *Yr32*. More recently, two new races emerged, *PstS13*c (year 2020), which gained virulence on *Yr27*, and the *PstS13*d (year 2021), which gained virulence on *Yr3* and *Yr25* in addition to the virulence pattern observed for *PstS13*b. The new *PstS13*d variant, virulent on both *Yr3* and *Yr25* could be the result of two subsequent single-set mutations. This hypothesis implies the existence of a race virulent on *Yr3* or *Yr25*, but it has not been detected in local surveys so far, which may be due to the small sample size or because it might have not been present in Uruguay. Another possible scenario of the appearance of *PstS13*d virulent on *Yr3* could be that this race variant was already present in the *PstS13* races, but it was not possible to determine with the differential lines used up to date.

The detection of *PstS13d* with virulence to *Yr3* and *Yr25* allows a better interpretation of the genes that might be present in some of the differential lines. Differential lines Vilmorin 23 and Nord Desprez, both carrying *Yr3* (Chen et al. 1996; Chen and Line 1993), resulted in a compatible interaction when tested with the *Yr25*-virulent race, which suggests that these differentials might also carry *Yr25*. The possible presence of *Yr25* in some differential lines, i.e., Carstens V, Spaldings Prolific and Strubes Dickkopf, was previously suggested (Calonnec and Johnson 1998; Eriksen et al. 2004; Calonnec et al. 2002). Following the assumption that differential lines carrying *Yr3* may also carry *Yr25*, indicate that *Yr3* is probably ineffective in previously reported *PstS13* races, which has been masked by a low IT conferred by the *Avr25/Yr25* phenotype. If the two *Yr3* differentials also carry *Yr25*, we cannot securely confirm virulence/avirulence for *Yr3* using the current differential lines, except for mutant isolates with virulence on *Yr25*. Interpretation of virulence entirely depends on knowledge about R-genes (known or unknown) in the differential lines, and many of the differential lines currently used could also contain other uncharacterized R-genes (Johnson 1992). In this context, the use of multiple differential lines representing each *Yr*-gene could be recommended for a more precise race typing analysis

In this study, although the number of samples was relatively low, we were able to detect three widely spread *Pst* genetic groups. However, additional *Pst* genetic groups could have been undetected. For example, *PstS14* reported in Argentina in 2017 (Carmona et al. 2019), was not detected in Uruguay despite the geographical proximity of both countries. *PstS14* was first detected in Northern Africa and Europe in 2016, causing severe epidemics on bread wheat in Morocco in 2017 (Hovmøller et al. 2018). Future surveys, with a larger number of samples collected at a regional scale, could complement this work and provide further insights into the understanding of the current distribution of genetic groups in the Southern Cone of South America.

## Conclusions

The hypothesis of initial migration into South America from distant source areas of *Pst* races responsible for current severe epidemics was confirmed. The identification of *Pst* races in Uruguay that were not previously described provides strong evidence of recent local evolution of *Pst*. The confirmation of the emergence of new races of *Pst* at the local level has implications for the management of the disease and plant breeding for disease resistance. In the short term, local management of ongoing epidemics on susceptible wheat varieties is limited to fungicide spraying. However, the deployment of resistant wheat cultivars appears as the most environmentally friendly strategy without additional cost for producers. The development of resistant cultivars implies permanent monitoring of the *Pst* races to generate cultivars with effective resistance to races present in a specific epidemiological zone. The occurrence of long-distance dispersal of races migrating from distant continents emphasizes the relevance of worldwide coordination of survey efforts. The new race variants reported in this work might migrate to other areas as well and potentially cause important yield losses in regions where cultivars that possess the overcome *Pst* resistance genes are deployed.

## Acknowledgment

The authors are grateful to Ellen Jørgensen, Janne Holm Hansen and Jakob Sørensen, Aarhus University, for technical assistance during *Pst* genotyping and phenotyping experiments at GRRC. We would also like to acknowledge Noelia Pérez for assistance in greenhouse tasks at the National Institute of Agricultural Research-INIA Uruguay.

