## Supplementary material for "New races with wider virulence indicate local evolution of *Puccinia striiformis* f. sp. *tritici* in South America"

- **Supplementary material 1**. Information of samples collected from spring-bread wheat in Uruguay during 2017-2021.

| Isolate ID | Region | Site | Latitude | Longitude | Altitude | Collection date |
|---|---|---|---|---|---|---|
| UY_147 | Soriano | Field | -33.596958 | -57.697885 | 105 | 07.11.2017 |
| UY_148 | Colonia | Field | -33.887676 | -58.365218 | 28 | 07.11.2017 |
| UY_149 | Rio Negro | Field | -33.077091 | -58.054353 | 25 | 07.11.2017 |
| UY_150 | Colonia | Trial | -34.336435 | -57.712592 | 72 | 26.10.2017 |
| UY_151 | Rio Negro | Trial | -32.687306 | -57.656250 | 70 | 26.10.2018 |
| UY_152 | Colonia | Trial | -34.336435 | -57.712592 | 72 | 29.10.2018 |
| UY_153 | Colonia | Trial | -34.336435 | -57.712592 | 72 | 08.11.2018 |
| UY_154 | Colonia | Trial | -34.336435 | -57.712592 | 72 | 08.11.2018 |
| UY_155 | Colonia | Trial | -34.336435 | -57.712592 | 72 | 08.11.2018 |
| UY_156 | Rio Negro | Trial | -32.679278 | -57.648778 | 79 | 08.11.2019 |
| UY_157 | Colonia | Trial | -34.336435 | -57.712592 | 72 | 23.10.2019 |
| UY_158 | Rio Negro | Trial | -32.683049 | -57.659589 | 87 | 14.10.2020 |
| UY_159 | Colonia | Trial | -34.336435 | -57.712592 | 72 | 19.10.2020 |
| UY_160 | Colonia | Trial | -34.336435 | -57.712592 | 72 | 11.11.2020 |
| UY_161 | Colonia | Trial | -34.336435 | -57.712592 | 72 | 10.11.2020 |
| UY_162 | Colonia | Trial | -34.336435 | -57.712592 | 72 | 10.11.2020 |
| UY_163 | Colonia | Trial | -34.336435 | -57.712592 | 72 | 10.11.2020 |
| UY_164 | Colonia | Trial | -34.336435 | -57.712592 | 72 | 10.11.2020 |
| UY_165 | Colonia | Trial | -34.336435 | -57.712592 | 72 | 10.11.2020 |
| UY_166 | Colonia | Trial | -34.336435 | -57.712592 | 72 | 10.11.2020 |
| UY_167 | Colonia | Trial | -34.336435 | -57.712592 | 72 | 23.11.2020 |
| UY_168 | Colonia | Trial | -34.336435 | -57.712592 | 72 | 23.11.2020 |
| UY_169 | Colonia | Trial | -34.339500 | -57.695393 | 65 | 28.10.2021 |
| UY_170 | Colonia | Trial | -34.339500 | -57.695393 | 65 | 28.10.2021 |
| UY_171 | Colonia | Trial | -34.339500 | -57.695393 | 65 | 28.10.2021 |
| UY_172 | Colonia | Trial | -34.339500 | -57.695393 | 65 | 28.10.2021 |
| UY_173 | Colonia | Trial | -34.339500 | -57.695393 | 65 | 28.10.2021 |

- **Supplementary material 2**. Allele sizes for each SSRs and genetic group assignment of samples collected in Uruguay during 2017-2021.

| | Multiplex-1 | | | | | | | | | | | | | | | | | | Multiplex-2 | | | | | | | | | | | | | | | | Single | | Single | | |
|---|---|---|---|---|---|---|---|---|---|---|---|---|---|---|---|---|---|---|---|---|---|---|---|---|---|---|---|---|---|---|---|---|---|---|---|---|---|---|---|
| Flourescence => | 6FAM | | 6FAM | | VIC | | VIC | | NED | | NED | | PET | | PET | | 6FAM | | 6FAM | | 6FAM | | VIC | | VIC | | NED | | NED | | PET | | PET | | PET | | NED | | |
| Motif size => | 3bp | | 3bp | | 3bp | | 3bp | | 3bp | | 2bp | | 2bp | | 3bp | | 3bp | | 3bp | | 3bp | | 2bp | | 3bp | | 2bp | | 2bp | | 2bp | | 3bp | | 2bp | | 2bp | | **Genetic** |
| **Isolate ID** | NRJO4 | | NRJO24 | | NRJN12 | | NRJN8 | | NRJN13 | | NRJN3 | | NRJN11 | | NRJN6 | | NRJO21 | | NRJN10 | | NRJO18 | | NWU-6 | | NRJO20 | | NRJN4 | | NRJN9 | | NRJN5 | | NWU-12 | | NRJO27 | | NRJN2 | | **group** |
| UY155 | 199 | 199 | 284 | 293 | 196 | 196 | 307 | 316 | 147 | 150 | 336 | 336 | 176 | 182 | 315 | 318 | 170 | 170 | 224 | 224 | 331 | 331 | 210 | 210 | 284 | 287 | 255 | 255 | 332 | 332 | 226 | 226 | 332 | 332 | 230 | 242 | nd | nd | PstS7 |
| UY163 | 199 | 199 | 284 | 293 | 196 | 196 | 307 | 316 | 147 | 150 | 336 | 336 | 176 | 182 | 315 | 318 | 170 | 170 | 224 | 224 | 331 | 331 | 210 | 210 | 284 | 287 | 255 | 255 | 332 | 332 | 226 | 226 | 332 | 332 | nd | nd | 169 | 169 | PstS7 |
| UY165 | 199 | 199 | 284 | 293 | 196 | 196 | 307 | 316 | 147 | 150 | 336 | 336 | 176 | 182 | 315 | 318 | 170 | 170 | 224 | 224 | 331 | 331 | 210 | 210 | 284 | 287 | 255 | 255 | 332 | 332 | 226 | 226 | 332 | 332 | 242 | 242 | nd | nd | PstS7 |
| UY173 | 199 | 199 | 284 | 293 | 196 | 196 | 307 | 316 | 147 | 150 | 336 | 336 | 176 | 182 | 315 | 318 | 170 | 170 | 224 | 224 | 331 | 331 | 210 | 210 | 284 | 287 | 255 | 255 | 332 | 332 | 226 | 226 | 332 | 332 | 230 | 230 | 169 | 169 | PstS7 |
| UY161 | 199 | 202 | 275 | 284 | 196 | 196 | 307 | 307 | 147 | 150 | 336 | 336 | 176 | 176 | 315 | 318 | 170 | 170 | 224 | 224 | 331 | 337 | 210 | 210 | 284 | 287 | 255 | 255 | 332 | 334 | 226 | 230 | 323 | 332 | 232 | 232 | 183 | 183 | PstS10 |
| UY166 | 199 | 202 | 275 | 284 | 196 | 196 | 307 | 307 | 147 | 150 | 336 | 336 | 176 | 176 | 315 | 318 | 170 | 170 | 224 | 224 | 331 | 337 | 210 | 210 | 284 | 287 | 255 | 255 | 332 | 334 | 226 | 230 | 323 | 332 | 232 | 232 | 183 | 183 | PstS10 |
| UY147 | 199 | 205 | 272 | 284 | 196 | 196 | 304 | 307 | 150 | 150 | 336 | 336 | 172 | 176 | 315 | 315 | 170 | 176 | 224 | 224 | 331 | 337 | 210 | 210 | 287 | 287 | 255 | 257 | 332 | 332 | 226 | 228 | 323 | 332 | 230 | 230 | 169 | 169 | PstS13 |
| UY148 | 199 | 205 | 272 | 284 | 196 | 196 | 304 | 307 | 150 | 150 | 336 | 336 | 172 | 176 | 315 | 315 | 170 | 176 | 224 | 224 | 331 | 337 | 210 | 210 | 287 | 287 | 255 | 257 | 332 | 332 | 226 | 228 | 323 | 332 | 230 | 230 | 169 | 169 | PstS13 |
| UY149 | 199 | 205 | 272 | 284 | 196 | 196 | 304 | 307 | 150 | 150 | 336 | 336 | 172 | 176 | 315 | 315 | 170 | 176 | 224 | 224 | 331 | 337 | 210 | 210 | 287 | 287 | 255 | 257 | 332 | 332 | 226 | 228 | 323 | 332 | nd | nd | 169 | 169 | PstS13 |
| UY150 | 199 | 205 | 272 | 284 | 196 | 196 | 304 | 307 | 150 | 150 | 336 | 336 | 172 | 176 | 315 | 315 | 170 | 176 | 224 | 224 | 331 | 337 | 210 | 210 | 287 | 287 | 255 | 257 | 332 | 332 | 226 | 228 | 323 | 332 | 230 | 230 | 169 | 169 | PstS13 |
| UY151 | 199 | 205 | 272 | 284 | 196 | 196 | 304 | 307 | 150 | 150 | 336 | 336 | 172 | 176 | 315 | 315 | 170 | 176 | 224 | 224 | 331 | 337 | 210 | 210 | 287 | 287 | 255 | 257 | 332 | 332 | 226 | 228 | 323 | 332 | 230 | 230 | 169 | 169 | PstS13 |
| UY152 | 199 | 205 | 272 | 284 | 196 | 196 | 304 | 307 | 150 | 150 | 336 | 336 | 172 | 176 | 315 | 315 | 170 | 176 | 224 | 224 | 331 | 337 | 210 | 210 | 287 | 287 | 255 | 257 | 332 | 332 | 226 | 228 | 323 | 332 | 230 | 230 | 169 | 169 | PstS13 |
| UY153 | 199 | 205 | 272 | 284 | 196 | 196 | 304 | 307 | 150 | 150 | 336 | 336 | 172 | 176 | 315 | 315 | 170 | 176 | 224 | 224 | 331 | 337 | 210 | 210 | 287 | 287 | 255 | 257 | 332 | 332 | 226 | 228 | 323 | 332 | 230 | 230 | 169 | 169 | PstS13 |
| UY154 | 199 | 205 | 272 | 284 | 196 | 196 | 304 | 307 | 150 | 150 | 336 | 336 | 172 | 176 | 315 | 315 | 170 | 176 | 224 | 224 | 331 | 337 | 210 | 210 | 287 | 287 | 255 | 257 | 332 | 332 | 226 | 228 | 323 | 332 | 230 | 230 | 169 | 169 | PstS13 |
| UY156 | 199 | 205 | 272 | 284 | 196 | 196 | 304 | 307 | 150 | 150 | 336 | 336 | 172 | 176 | 315 | 315 | 170 | 176 | 224 | 224 | 331 | 337 | 210 | 210 | 287 | 287 | 255 | 257 | 332 | 332 | 226 | 228 | 323 | 332 | 230 | 230 | 169 | 169 | PstS13 |
| UY157 | 199 | 205 | 272 | 284 | 196 | 196 | 304 | 307 | 150 | 150 | 336 | 336 | 172 | 176 | 315 | 315 | 170 | 176 | 224 | 224 | 331 | 337 | 210 | 210 | 287 | 287 | 255 | 257 | 332 | 332 | 226 | 228 | 323 | 332 | 230 | 230 | 169 | 169 | PstS13 |
| UY158 | 199 | 205 | 272 | 284 | 196 | 196 | 304 | 307 | 150 | 150 | 336 | 336 | 172 | 176 | 315 | 315 | 170 | 176 | 224 | 224 | 331 | 337 | 210 | 210 | 287 | 287 | 255 | 257 | 332 | 332 | 226 | 228 | 323 | 332 | 230 | 230 | 169 | 169 | PstS13 |
| UY159 | 199 | 205 | 272 | 284 | 196 | 196 | 304 | 307 | 150 | 150 | 336 | 336 | 172 | 176 | 315 | 315 | 170 | 176 | 224 | 224 | 331 | 337 | 210 | 210 | 287 | 287 | 255 | 257 | 332 | 332 | 226 | 228 | 323 | 332 | 230 | 230 | 169 | 169 | PstS13 |
| UY160 | 199 | 205 | 272 | 284 | 196 | 196 | 304 | 307 | 150 | 150 | 336 | 336 | 172 | 176 | 315 | 315 | 170 | 176 | 224 | 224 | 331 | 337 | 210 | 210 | 287 | 287 | 255 | 257 | 332 | 332 | 226 | 228 | 323 | 332 | 230 | 230 | 169 | 169 | PstS13 |
| UY167 | 199 | 205 | 272 | 284 | 196 | 196 | 304 | 307 | 150 | 150 | 336 | 336 | 172 | 176 | 315 | 315 | 170 | 176 | 224 | 224 | 331 | 337 | 210 | 210 | 287 | 287 | 255 | 257 | 332 | 332 | 226 | 228 | 323 | 332 | 230 | 230 | 169 | 169 | PstS13 |
| UY168 | 199 | 205 | 272 | 284 | 196 | 196 | 304 | 307 | 150 | 150 | 336 | 336 | 172 | 176 | 315 | 315 | 170 | 176 | 224 | 224 | 331 | 337 | 210 | 210 | 287 | 287 | 255 | 257 | 332 | 332 | 226 | 228 | 323 | 332 | 230 | 230 | 169 | 169 | PstS13 |
| UY170 | 199 | 205 | 272 | 284 | 196 | 196 | 304 | 307 | 150 | 150 | 336 | 336 | 172 | 176 | 315 | 315 | 170 | 176 | 224 | 224 | 331 | 337 | 210 | 210 | 287 | 287 | 255 | 257 | 332 | 332 | 226 | 228 | 323 | 332 | 230 | 230 | 169 | 169 | PstS13 |
| UY171 | 199 | 205 | 272 | 284 | 196 | 196 | 304 | 307 | 150 | 150 | 336 | 336 | 172 | 176 | 315 | 315 | 170 | 176 | 224 | 224 | 331 | 337 | 210 | 210 | 287 | 287 | 255 | 257 | 332 | 332 | 226 | 228 | 323 | 332 | 230 | 230 | 169 | 169 | PstS13 |
| UY172 | 199 | 205 | 272 | 284 | 196 | 196 | 304 | 307 | 150 | 150 | 336 | 336 | 172 | 176 | 315 | 315 | 170 | 176 | 224 | 224 | 331 | 337 | 210 | 210 | 287 | 287 | 255 | 257 | 332 | 332 | 226 | 228 | 323 | 332 | 230 | 230 | 169 | 169 | PstS13 |
| UY162 | 199 | 205 | 272 | 284 | 196 | 196 | 304 | 307 | 150 | 150 | 336 | 336 | 172 | 176 | 315 | 315 | 170 | 176 | 224 | 224 | 331 | 337 | 210 | 210 | 287 | 287 | 255 | 257 | 332 | 332 | 226 | 228 | 323 | 332 | 230 | 230 | 169 | 169 | PstS13 |
| UY164 | 199 | 205 | 272 | 284 | 196 | 196 | 304 | 307 | 150 | 150 | 336 | 336 | 172 | 176 | 315 | 315 | 170 | 176 | 224 | 224 | 331 | 337 | 210 | 210 | 287 | 287 | 255 | 257 | 332 | 332 | 226 | 228 | 323 | 332 | 230 | 230 | 169 | 169 | PstS13 |
| UY169 | 199 | 205 | 272 | 284 | 196 | 196 | 304 | 307 | 150 | 150 | 336 | 336 | 172 | 176 | 315 | 315 | 170 | 176 | 224 | 224 | 331 | 337 | 210 | 210 | 287 | 287 | 255 | 257 | 332 | 332 | 226 | 228 | 323 | 332 | 230 | 230 | 169 | 169 | PstS13 |

- **Supplementary material 3**. Infection types on *Pst* differential lines of *Pst* isolates collected in Uruguay during 2017-2021.

| Variety => | Chinese 166 | Kalyansona | Vilmorin 23 | Hybrid 46 | Suwon | Heines Kolben | Avocet *Yr6* | Lee | Avocet *Yr8* | Avocet *Yr9* | Moro | Cortez | VPM1 | Avocet *Yr17* | TP 981 | Opata | Avocet *Yr32* | Spaldings Prolific | Avocet *S* | Ambition | Benchmark | Kalmar | KWS Zyatt | Nemo | Virulence phenotype | Race |
|---|---|---|---|---|---|---|---|---|---|---|---|---|---|---|---|---|---|---|---|---|---|---|---|---|---|---|
| Yr-genes => | *1* | *2,+* | *3,+* | *4,+* | *Su* | *6,+* | *AvS,6* | *7,+* | *AvS,8* | *AvS,9* | *10* | *15* | *17,+* | *AvS,17* | *25,+* | *18,27,+* | *AvS,32* | *Sp,25,+* | *AvS* | *Amb* | | | | | | |
| UY155 | 8 | 8 | 6,7 | 5,6,7 | 8 | 8 | 8 | 8 | 1 | 8 | 1 | 1 | 7 | 7 | 7 | 3,4 | 7 | 7 | 8 | 7 | 7 | 4,5,6 | 7 | 3,4,5 | 1,2,3,4,6,7,-,9,-,-,17,25,-,32,Sp,AvS,Amb | Warrior |
| UY163 | 8 | 8 | 6,7 | 5,6 | 8 | 8 | 7 | 8 | 1 | 8 | 1 | 1 | 7,8 | 8 | 7 | 3,4 | 8 | 7 | 8 | 8 | 6,7 | 3,4,5 | 7 | 4,5,6 | 1,2,3,4,6,7,-,9,-,-,17,25,-,32,Sp,AvS,Amb | Warrior |
| UY165 | 8 | 7 | 6,7 | 5,6 | 8 | 8 | 8 | 8 | 0 | 8 | 1 | 1 | 7 | 7 | 8 | 3 | 7 | 7 | 8 | 8 | 8 | 4,5,6 | 7 | 4,5,6 | 1,2,3,4,6,7,-,9,-,-,17,25,-,32,Sp,AvS,Amb | Warrior |
| UY173 | 7 | 7 | 7 | 5,6 | 8 | 8 | 8 | 8 | 1 | 8 | 1 | 1 | 7 | 7 | 7 | 3,4 | 8 | 7 | 8 | 7 | 7 | 5,6,7 | 7 | 4,5 | 1,2,3,4,6,7,-,9,-,-,17,25,-,32,Sp,AvS,Amb | Warrior |
| UY161 | 8 | 8 | 8 | 5,6 | 8 | 8 | 8 | 8 | 2,3 | 8 | 2,3 | 1 | 8 | 8 | 7 | nd | 7 | 6,7 | 8 | 3,4 | 7 | 3,4,5 | 6,7 | 2,3 | 1,2,3,4,6,7,-,9,-,-,17,25,-,32,Sp,AvS,- | Benchmark |
| UY166 | 7 | 8 | 6,7 | 5 | 7 | 7 | 7 | 7 | 1 | 8 | 1 | 1 | 6,7 | 7 | 7 | 3,4 | 7 | 7 | 7 | nd | 7 | 4,5,6,7 | 6,7 | 3,4,5 | 1,2,3,4,6,7,-,9,-,-,17,25,-,32,Sp,AvS,- | Benchmark |
| UY147 | 1 | 8 | 1 | 4,5 | 8 | 8 | 7 | 8 | 8 | 8 | 2,3 | 0,1 | 1 | 2,3 | 3,4,3 | 4 | 2,3 | 5,6,7 | 8 | 3,4 | 0,1 | 0,1 | 1 | 1,2 | -,2,-,-,6,7,8,9,-,-,-,-,-,-,-,AvS,- | Triticale2015 |
| UY148 | 1 | 7 | 1 | 2,3,4 | 7 | 8 | 7 | 8 | 8 | 7 | 2,3 | 1,2 | 1,2 | 2,3 | 3,4 | 4,5,6 | 2,3 | 5,6,7 | 7 | 4,5 | 1 | 0,1 | 1 | 2 | -,2,-,-,6,7,8,9,-,-,-,-,-,-,-,AvS,- | Triticale2015 |
| UY149 | 1 | 8 | 1,2 | 3,4 | 8 | 8 | 7 | 8 | 8 | 8 | 2,3 | 1 | 1 | 2,3 | 3,4 | 5,6 | 3,4 | 5,6,7 | 7 | 4,5,6 | 0,1 | 0,1 | 0 | 1,2 | -,2,-,-,6,7,8,9,-,-,-,-,-,-,-,AvS,- | Triticale2015 |
| UY150 | 1 | 8 | 1,2 | 3,4 | 7 | 8 | 8 | 8 | 8 | 8 | 2 | 1,2 | 1 | 2,3 | 3,4,5 | 4 | 3,4 | 5,6,7 | 8 | 4,5 | 2 | 1 | 1 | 1 | -,2,-,-,6,7,8,9,-,-,-,-,-,-,-,AvS,- | Triticale2015 |
| UY151 | 1 | 8 | 1 | 3,4 | 8 | 8 | 7 | 8 | 8 | 7,8 | 2,3 | 1 | 1,2 | 2,3 | 3 | 3,4,5 | 4 | 6,7 | 7 | 5,6 | 0,1 | 0,1 | 1 | 1,2 | -,2,-,-,6,7,8,9,-,-,-,-,-,-,-,AvS,- | Triticale2015 |
| UY152 | 1 | 8 | 1,2 | 4 | 8 | 8 | 8 | 8 | 8 | 8 | 3 | 1 | 1,2 | 1,2 | 3,4 | 4 | 2,3,4 | 5,6,7 | 8 | 5,6 | 1 | 1 | 1 | 1,2 | -,2,-,-,6,7,8,9,-,-,-,-,-,-,-,AvS,- | Triticale2015 |
| UY153 | 1 | 8 | 1 | 3,4,5 | 8 | 8 | 8 | 8 | 8 | 8 | 2,3 | 0,1 | 1 | 1,2 | 3 | 3,4,5 | 4,5 | 5,6,7 | 8 | 5,6,7 | 1 | 1 | 1,2 | 1,2 | -,2,-,-,6,7,8,9,-,-,-,-,-,-,-,AvS,- | Triticale2015 |
| UY154 | 1 | 8 | 2,3 | 3,4 | 8 | 8 | 8 | 8 | 8 | 8 | 2,3 | 1,2 | 1,2 | 1,2 | 3,4 | 3,4,5 | 4 | 5,6,7 | 8 | 4,5,6 | 1 | 0,1 | 1,2 | 1 | -,2,-,-,6,7,8,9,-,-,-,-,-,-,-,AvS,- | Triticale2015 |
| UY156 | 1 | 8 | 2 | 3,4 | 7,8 | 8 | 8 | 8 | 8 | 8 | 2 | 1 | 1,2 | 8 | 3,4 | 5,6 | 8 | 5,6,7 | 8 | 4,5 | 1 | 1 | 1,2 | 1,2 | -,2,-,-,6,7,8,9,-,-,17,-,-,32,-,AvS,- | Triticale2015 |
| UY157 | 1 | 8 | 1 | 3,4 | 8 | 8 | 8 | 8 | 8 | 8 | 3,4,5 | 1 | 1,2 | 7 | 3,4 | 4,5 | 7 | 5,6,7 | 8 | 5,6 | 1 | 1 | 1 | 1,2 | -,2,-,-,6,7,8,9,-,-,17,-,-,32,-,AvS,- | Triticale2015 |
| UY158 | 1 | 8 | 2 | 4,5 | 7 | 8 | 8 | 8 | 8 | 8 | 2 | 2 | 2 | 7,8 | 2 | 4,5 | 8 | 5,6,7 | 8 | 5,6 | 1 | 1 | nd | 1 | -,2,-,-,6,7,8,9,-,-,17,-,-,32,-,AvS,- | Triticale2015 |
| UY159 | 1 | 8 | 1,2 | 3,4 | 7 | 8 | 8 | 8 | 8 | 8 | 2 | 1,2 | 1,2,3 | 7,8 | 4,5 | 6 | 8 | 5,6,7 | 8 | 5,6 | 1 | 1 | 1 | 1,2 | -,2,-,-,6,7,8,9,-,-,17,-,-,32,-,AvS,- | Triticale2015 |
| UY160 | 1 | 8 | 1 | 3,4 | 7 | 7 | 8 | 8 | 7 | 8 | 2 | 1,2 | 1 | 8 | 5 | 6 | 8 | 5,6,7 | 8 | 5,6 | 1,2 | 1 | 1,2 | 1,2 | -,2,-,-,6,7,8,9,-,-,17,-,-,32,-,AvS,- | Triticale2015 |
| UY162 | 1 | 7 | 1 | 4,5 | 7 | 8 | 7 | 8 | 8 | 8 | 2,3 | 1 | 1,2 | 7 | 3,4 | 5 | 7 | 5,6,7 | 8 | 4,5,6 | 0,1 | 0,1 | 0,1 | 1,2 | -,2,-,-,6,7,8,9,-,-,17,-,-,32,-,AvS,- | Triticale2015 |
| UY164 | 1 | 8 | 1 | 3,4 | 8 | 8 | 8 | 8 | 8 | 8 | 2,3 | 1 | 1,2 | 7 | 3,4 | 6 | 8 | 5,6,7 | 8 | 4,5 | 1 | 0,1 | 1 | 1,2 | -,2,-,-,6,7,8,9,-,-,17,-,-,32,-,AvS,- | Triticale2015 |
| UY167 | 1 | 8 | 1,2 | 3,4 | 7,8 | 8 | 8 | 8 | 8 | 8 | 2,3 | 2 | 1,2 | 8 | 3,4 | 5 | 8 | 5,6,7 | 8 | 5,6 | 0,1 | 0,1 | 1 | 1,2 | -,2,-,-,6,7,8,9,-,-,17,-,-,32,-,AvS,- | Triticale2015 |
| UY170 | 1 | 8 | 1 | 3,4 | 7 | 7 | 7 | 7 | 7 | 7 | 2,3 | 1 | 1 | 7 | 2,3 | 6 | 7 | 5,6,7 | 7 | 4,5 | 1 | 0,1 | 1 | 2 | -,2,-,-,6,7,8,9,-,-,17,-,-,32,-,AvS,- | Triticale2015 |
| UY171 | 1 | 8 | 1,2 | 4 | 8 | 8 | 8 | 8 | 8 | 8 | 2,3,4 | 1 | 1 | 7 | 3 | 5 | 7 | 5,6,7 | 7 | 5,6 | 0,1 | 0,1 | 1 | 1,2 | -,2,-,-,6,7,8,9,-,-,17,-,-,32,-,AvS,- | Triticale2015 |
| UY172 | 1 | 8 | 1 | 2,3,4 | 8 | 8 | 8 | 8 | 8 | 8 | 2,3 | 1,2 | 1,2 | 7 | 2,3 | 5,6 | 7 | 5,6,7 | 7 | 4,5 | 0,1 | 0,1 | 1 | 1 | -,2,-,-,6,7,8,9,-,-,17,-,-,32,-,AvS,- | Triticale2015 |
| UY168 | 1 | 8 | 1,2 | 2,3,4 | 7,8 | 7 | 8 | 8 | 8 | 8 | 3 | 1 | 1,2 | 7 | 3,4 | 7 | 7,8 | 6 | 8 | 5,6 | 1 | 0,1 | 1 | 1,2 | -,2,-,-,6,7,8,9,-,-,17,-,27,32,-,AvS,- | Triticale2015 |

Note. Race typing was performed using a 0–9 infection type scale (McNeal et al. 1971), scores between 7 and 9 indicate compatibility (virulence), and scores equal to or below 6 indicate incompatibility (avirulence).

- **Supplementary material 4**. Infection types on *Pst* differential lines of *Pst13* variants tested in Supplementary test.

| Variety => | Avocet *S* | Strubes Dickkopf | Spaldings Prolific | Kalyansona | Heines Peko | Heines VII | Nord Desprez | Vilmorin 23 | VPM1 | TP 981 | Avocet *Yr17* | Carstens V | Avocet *Yr32* | Avocet *Yr27* | Opata | Race |
|---|---|---|---|---|---|---|---|---|---|---|---|---|---|---|---|---|
| *Yr*-genes => | *AvS* | *Sd,25* | *Sp,25,+* | *2,+* | *2,6,25,+* | *2,25,+* | *3,+* | *3,+* | *17,+* | *25,+* | *AvS,17* | *25,32,+* | *AvS,32* | *AvS,27* | *18,27,+* | |
| DK69 | 7 | 1 | 5,6,7 | 7 | 2 | 1 | 2 | 1 | 0,1 | 2 | 0 | 0,1 | 0,2,3 | 0 | 2,3,4 | Triticale 2015 a |
| UY147 | 7 | 1 | 5,6 | 7 | 2 | 1 | 2 | 1 | 1 | 2,3,4 | 0 | 0,1 | 2,3 | 0 | 1,2,3 | Triticale 2015 a |
| UY150 | 7 | 1 | 5,6,7 | 7 | 2 | 1 | 2 | 1 | 1 | 2,3,4 | 0,1,2 | 0,1,2 | 2,3 | 0,1 | 0,1,2 | Triticale 2015 a |
| UY152 | 7 | 1 | 5,6,7 | 7 | 2 | 1 | 2 | 1 | 0,1 | 2,3,4 | 0,1,2 | 0,1,2 | 2,3 | 0,1 | 0,1,2 | Triticale 2015 a |
| UY158 | 7 | 1 | 5,6,7 | 7 | 2 | 1 | 2 | 1 | 0,1 | 2 | 7 | 0,1 | 7 | 3,4 | 4,5 | Triticale 2015 b |
| UY168 | 7 | 1 | 5,6,7 | 7 | 2 | 1 | 2 | 1 | 0,1 | 2 | 6,7 | 0,1,2 | 7 | 7 | 7 | Triticale 2015 c |
| UY623 | 7 | 2,3 | 6,7 | 7 | 7 | 5,6 | 7 | 5,6 | 2,3 | 7 | 7 | 6,7 | 7 | 4,5,6 | 4,5 | Triticale 2015 d |

Note. The first and the second leaves were scored using a 0–9 scale (McNeal et al. 1971), scores between 7 and 9 indicate compatibility (virulence), and scores equal to or below 6 indicate incompatibility (avirulence). Letters a-d indicate different virulence phenotypes within genetic group *PstS13*.
